# RNAi screen in *Tribolium* reveals involvement of F-BAR proteins in myoblast fusion and visceral muscle morphogenesis in insects

**DOI:** 10.1101/397232

**Authors:** Dorothea Schultheis, Jonas Schwirz, Manfred Frasch

## Abstract

In a large-scale RNAi screen in *Tribolium castaneum* for genes with knock-down phenotypes in the larval somatic musculature, one recurring phenotype was the appearance of larval muscle fibers that were significantly thinner than those in control animals. Several of the genes producing this knock-down phenotype corresponded to orthologs of *Drosophila* genes that are known to participate in myoblast fusion, particularly via their effects on actin polymerization. A new gene previously not implicated in myoblast fusion but displaying a similar thin-muscle knock-down phenotype was the *Tribolium* ortholog of *Nostrin*, which encodes an F-BAR and SH3 domain protein. Our genetic studies of *Nostrin* and *Cip4*, a gene encoding a structurally related protein, in *Drosophila* show that the encoded F-BAR proteins jointly contribute to efficient myoblast fusion during larval muscle development. Together with the F-Bar protein Syndapin they are also required for normal embryonic midgut morphogenesis. In addition, *Cip4* is required together with *Nostrin* during the profound remodeling of the midgut visceral musculature during metamorphosis. We propose that these F-Bar proteins help govern proper morphogenesis particularly of the longitudinal midgut muscles during metamorphosis.

## Introduction

As described in the accompanying paper (Schultheis *et al.* 2019), we participated in large-scale screens with systemic RNAi in the flour beetle *Tribolium castaneum* aiming to identify new genes that regulate the development of the somatic musculature. One screen was for knock-down phenotypes in muscles of late stage embryos and first instar larvae, which involved injecting double stranded RNAs into pupae of a tester strain that expressed EGFP in all somatic (and visceral) muscles. A second screen was for knockdown phenotypes in the adult indirect flight muscles of the thorax of late stage pupae, which involved injections into larvae of a strain expressing EGFP in these muscles. A broad overview over these screens, which included screening for various other phenotypes, has been presented in Schmitt-Engel *et al.* (2015). After identifying new genes associated with knock-down phenotypes in the somatic musculature in *Tribolium* our main strategy was to utilize the superior genetic tools and accrued body of information in *Drosophila* to study the functions of their fly orthologs in detail and place them into the known regulatory framework of muscle development in the fly.

Herein we focus on genes that we selected based on their larval muscle phenotypes in the pupal injection screen. Specifically, this is a group of genes that produced a phenotype of somatic muscles in embryos that were significantly thinner as compared to controls, which led to anomalous gaps between parallel muscle fibers. The *Drosophila* orthologs of several of these genes are known to participate in myoblast fusion during embryonic muscle development in the fly, particularly via their effects on promoting actin polymerization.

*Drosophila* myoblast fusion is an increasingly well-characterized process, during which a set number of fusion-competent myoblasts fuses with a single muscle founder cell and with the nascent myotube formed by this process. The asymmetry of this process relies on the cell type specific expression of several of the key components of the recognition and fusion machinery (Kim *et al.* 2015; Deng *et al.* 2017). In particular, the recognition and adhesion of the two types of myoblast involves the engagement of the immunoglobulin domain proteins Sticks-and-stones (Sns) and Hibris (Hbs) on the surface of the fusion-competent myoblasts with the structurally related proteins Kin of irre (Kirre) (aka, Dumbfounded, Duf) and Roughest (Rst, aka, IrreC) on the surface of the muscle founder cells. This interaction then triggers downstream events in both cell types, which culminate in the differential assembly of polymerized actin structures at the prospective fusion site in fusion-competent versus founder myoblasts. Membrane breakdown and fusion pores occur upon the extension of actin-propelled protrusions from the fusion-competent myoblasts that invade the founder cells, and of F-actin sheaths thought to act as counter-bearings underneath the opposing membranes of the founder cells. The concomitant assembly of ring-shaped multiprotein complexes and the removal of cell adhesion proteins such as N-Cadherin at these sites additionally promote and orchestrate the formation and extension of fusion pores at these sites (Önel and Renkawitz-Pohl 2009; Önel *et al.* 2014). Whether any fusogens, as known to be active in other contexts of cell fusion (Segev *et al.* 2018), are involved in membrane fusions in *Drosophila* myoblast fusion is currently not known. Consecutive rounds of myoblast fusions generate the multinucleated muscle precursors in this manner.

A new gene identified based on its thin-muscle phenotype in *Tribolium castaneum (Tc)* was *TC013784 (Tc-Nostrin)*, homologs of which previously have not been implicated in *Drosophila* myoblast fusion. This gene encodes a protein with an F-BAR domain within its N-terminal half and an SH3 domain at its C-terminus. F-BAR proteins associate as curved homo-dimers with the inner face of the plasma membrane via binding to phospholipids and regulate membrane curvature as well as actin polymerization in various contexts (Roberts-Galbraith and Gould 2010; Liu *et al.* 2015; Salzer *et al.* 2017). Here we focus on the analysis of *Drosophila Nostrin* and two related genes, *Cip4* and *Syndapin (Synd)*, within this superfamily that encode F-BAR plus SH3 domain proteins, during muscle development. Previously, the functions of these *Drosophila* genes have been characterized within other developmental contexts, including germ line cell encapsulation *(Nost*; Zobel *et al.* 2015), the formation of proper numbers of wing hairs *(Cip4;* Fricke *et al.* 2009), and postsynaptic membrane organization *(Synd*; Kumar *et al.* 2009b). As described herein, our genetic analysis of these F-BAR genes in *Drosophila* muscle development shows that their encoded proteins, particularly Nostrin and Cip4, make joint contributions to myoblast fusion during embryogenesis. We also show that these F-BAR proteins, with a predominant role of Cip4, are critical for normal morphogenesis of the adult visceral muscles, which undergo major remodeling processes during metamorphosis.

## Materials and Methods

### *Tribolium* iBeetle database

RNAi phenotypes from the screen and additional gene information can be can be looked up in the iBeetle database (http://ibeetle-base.uni-goettingen.de). There is only one representative each for *Nost* and *Synd* in the *Tribolium* genome, but there are two representatives for *Cip4, TC014985* and *TC034900.* Other Bar domain genes are more distantly related such as *TC005182 (Tc-CG8176)* (AH/Bar domain class). Except for *Nost*, none of these others were denoted with a highly penetrant or specific muscle phenotype, which in the case of *Cip4* could be due to functional redundancy of the paralogs.

### *Drosophila* strains

All Drosophila melanogaster stocks were kept on standard medium at 25°C. The following *Drosophila* strains were used in this study: *Cip4^Δ32^* (Fricke *et al.* 2009); *Cip4^Δ32^,Synd^1d^/TM3, twi>>GFP* (this work); *Cip4^Δ32^, Synd^dEx22^/TM3, twi>>GFP* (this work); *Cip4::YFP (PBac{754.P.FSVS-0}Cip4*^CPTI03231^, Kyoto Stock Center); *hsFLP/TM6* (Bloomington Drosophila Stock Center, #279); *lmd^1^/TM3, Kr>>lacZ* (Duan *et al.* 2001); *UAS-Nostrin-EGFP* (Zobel *et al.* 2015); *Nost^df004^* (this work); *Nos^df016^* (this work); *Nost^df004^*;;*Cip4 ^Δ32^,Synd^1d^/TM3, twi>>GFP* (this work); *Nost^df004^;;Cip4*^Δ32^, *Synd^dEx22^/TM3, twi>>GFP* (this work); *Nost^df004^;;Cip4*^Δ32^/*TM3,hs-hid3* (Zobel *et al.* 2015)(and this work); *Nost^df004^;;Cip4^Δ32^/TM3, twi>>GFP* (this work); *Nost^df004^;;Synd^1d^/TM3, twi>>GFP* (this work); *Nost^df004^;;Synd^dEx22^/TM3,twi>>GFP* (this work); *P{XP}CG10962^d08142^* (Harvard Medical School); *PBac{WH}CG10962f^02373^* (Harvard Medical School); *PBac{WH}f06363* (Harvard Medical School); *rP289-lacZ* (Nose *et al.* 1998); *Synd^1d^/TM3,twi>>GFP* (Kumar *et al.* 2009b); *Synd^dEx22^/TM3,twi>>GFP* (Kumar *et al.* 2009a); *Mef2-GAL4* (Ranganayakulu *et al.* 1998).

### Generation of *Nostrin* mutants

*Nostrin* mutants were generated via the *flp*/FRT-system as described in Parks *et al.* (2004). For the generation of *Nostrin* deletions either the strain *P{XP}CG10962^d08142^* or the strain *PBac{WH}CG10962^f02373^* in combination with *PBac{WH}f06363* were used (Figure S2). The deletions were identified via PCR. The resulting *Nostrin* deletion mutant strains *Nost^df004^* and *Nost^ff016^* are fully viable and fertile.

### Generation of homozygous *Nostrin* and *Cip4* and of *Cip4 Syndapin* double mutants

To obtain adult flies homozygous mutant for both *Nostrin* and *Cip4* (lacking the zygotic and maternal expression of both genes) the strain *Nost^df004^;;Cip4^Δ32^/TM3,hs-hid3* (see also Zobel *et al.* 2015) was used. Heat shocking the progeny for 1,5 hours at 37°C during 3 – 4 days resulted in the survival of only homozygous double mutant escaper flies and the death of all animals carrying the balancer chromosome. These escapers were mated with each other for obtaining *Nost;;Cip4* (m+z) embryos and (rare) adults for analyzing the embryonic somatic and adult midgut musculatures.

Meiotic recombinants carrying both *Cip4* and *Synd* on chromosome 3 were identified by checking for lethality (due to *Synd*) along with the presence of wing hair duplications (not present in *Synd* single mutants).

### Staining procedures

*Drosophila* embryo fixations, immunostainings for proteins and RNA *in situ* hybridization were performed as described previously (Azpiazu and Frasch 1993; Knirr *et al.* 1999). The Elite ABC-HRP kit (Vector Laboratories) and TSA Cyanine 3 System and TSA Fluorescein System (PerkinElmer Inc.) were used for fluorescent detection of RNA and One-Step NBT/BCIP (Thermo Scientific) for the non-fluorescent detection of RNA. The following antibodies were used: sheep anti-Digoxigenin (1:2000; Roche), sheep anti-Digoxigenin conjugated with alkaline phosphatase (1:2000; Roche), mouse antiEven-Skipped (1:100; DSHB, Iowa), mouse anti-GFP (1:100; Invitrogen), rabbit anti-GFP (1:2000; Invitrogen), rabbit anti-Mef2 (1:750) (Bour *et al.* 1995), rat anti-Org-1 (1:100) (Schaub *et al.* 2012), rabbit anti-Tinman (1:750) (Yin *et al.* 1997), rat anti-Tropomyosin1 (1:200; Babraham Institute), rabbit anti-β3Tubulin (1:3000; gift from R. Renkawitz-Pohl), rabbit anti-β-Galactosidase (1:1500; Promega), mouse anti-β-Galactosidase (40-1a) (1:20; DSHB, Iowa), mouse anti-lamin (T40) (1:25; Frasch *et al.* 1988) and digoxigenin labeled *nostrin, cip4* and *syndapin* antisense RNA probes (all 1:200). Secondary antibodies used were conjugated with DyLight 488, DyLight 647 or DyLight 549 (1:200; Jackson Immuno Research) or with biotin (1:500; Dianova). The following digoxigenin-labeled RNA antisense probes were used: *Nostrin, Cip4* and *Syndapin.* T7 promotor-tagged templates were generated by PCR (for primers see supplement) from cDNA clones obtained from the *Drosophila Genomics Resource Center (Nostrin:* clone #IP202041; *Cip4*: clone #FI02049; *Syndapin*: clone #LD46328). *Tribolium* fixation and *in situ* hybridization were performed as described previously (Tautz and Pfeifle 1989; Patel *et al.* 1994). For the generation of RNA antisense probes the same primers as for the dsRNA fragments were used (see http://ibeetle-base.uni-goettingen.de/gb2/gbrowse/tribolium/). Images were acquired on a Leica SP5II confocal laser scanning microscope using a HC PL APO20x/0.70 and HCX PL APO 63x/1.3 objectives (with glycerol) and the LAS AF (Leica) software, on an Axio Imager (Zeiss) equipped with an ApoTome (Zeiss) and Plan Apochromat 20x/0.8, 40x/1.3, 63x/1.4 objectives using the Axiovision4.8 software or on an Axio Scope A1 (Zeiss) using the ProgRes CapturePRo (Jenopik) software. The final figures were obtained using Photoshop CS5 (Adobe).

### Analysis of adult gut phenotypes and wing bristle phenotypes

Adult flies were narcotized with CO_2_. After cutting off the head, the flies were pinned through the thorax with their ventral side facing up onto a wax dish. After covering the flies with PBT the abdomen was opened along the ventral side. Next the gut was removed from the abdomen using forceps, transferred into a staining dish, and fixed for 20 – 40 min in PBS containing 3,7% formaldehyde. Guts were stained over night at 4°C with Phalloidin-Atto-550 (1:3000; Sigma-Aldrich), washed three times with PBS and embedded in Vectashield (Vector Laboratories).

For the analysis of the wing bristles adult flies were narcotized with CO_2_, the wings were removed at the notum using forceps and embedded in Euparal (Roth).

### Research materials and data availability

Materials produced in this study are available upon request. The authors affirm that all data necessary for confirming the conclusions of this article are represented fully within the article and its tables and figures with the exception of sequence information (e.g., for amplification primers) that is available at http://ibeetle-base.uni-goettingen.de/gb2/gbrowse/tribolium/.

## Results

### Knock-downs of orthologs of *Drosophila* genes involved in myoblast fusion cause ‘thin-muscle’ phenotypes

When we inspected the muscle phenotypes of genes for which their *Drosophila* orthologs have been implicated in myoblast fusion in the iBeetle database, we noticed that in many (albeit not all) cases these displayed significantly thinner muscles after their knock-down (Fig. 1A – F). This phenotype is particularly obvious for the dorsal and ventral longitudinal muscles, which normally are broad and touch their neighbors aligned in parallel (Fig. 1A). By contrast, upon knock-down of the *Tribolium* orthologs of *Drosophila ced-12* (aka *Elmo), Crk oncogene (Crk), schizo (siz)* (aka *loner)* and *Verprolin1 (Vrp1)* (aka *solitary, sltr)*, all of which are known to participate in myoblast fusion, these muscles are thinner and therefore clear gaps are present between them (Fig. 1B – E) (Chen *et al.* 2003; Balagopalan *et al.* 2006; Kim *et al.* 2007; Massarwa *et al.* 2007; Geisbrecht *et al.* 2008; Jin *et al.* 2011). Similar effects are seen upon knock-down of the *Tribolium* ortholog of *lameduck (lmd)*, which in *Drosophila* is needed for specifying fusion-competent myoblasts (Fig. 1F) (Duan *et al.* 2001). Of note, this phenotype differs from the prototypical myoblast fusion phenotype in *Drosophila*, which is characterized by the presence of large numbers of unfused myoblasts. However, also in *Drosophila* fusion mutants the unfused myoblasts tend to disappear at late embryonic stages, presumably because of cell death. In *Drosophila* mutants for genes with less prominent functions, with weak alleles, or with partial functional rescue by maternal products, the muscles are thinner as well due to the reduced uptake of fusion competent cells (e.g., Hamp *et al.* 2016). We propose that the ‘thin muscle’ phenotypes in *Tribolium* knockdowns of most myoblast fusion genes (including some weak phenotypes with *Tcas kirre/rst;* Schultheis *et al.* 2019) result from similar effects of incomplete functional knock-down and rapid disappearance of the unfused myoblasts. The absence of the GFP marker at earlier stages unfortunately prevented the detection of unfused myoblasts in control and RNAi treated embryos to confirm this explanation.

**Figure 1.**
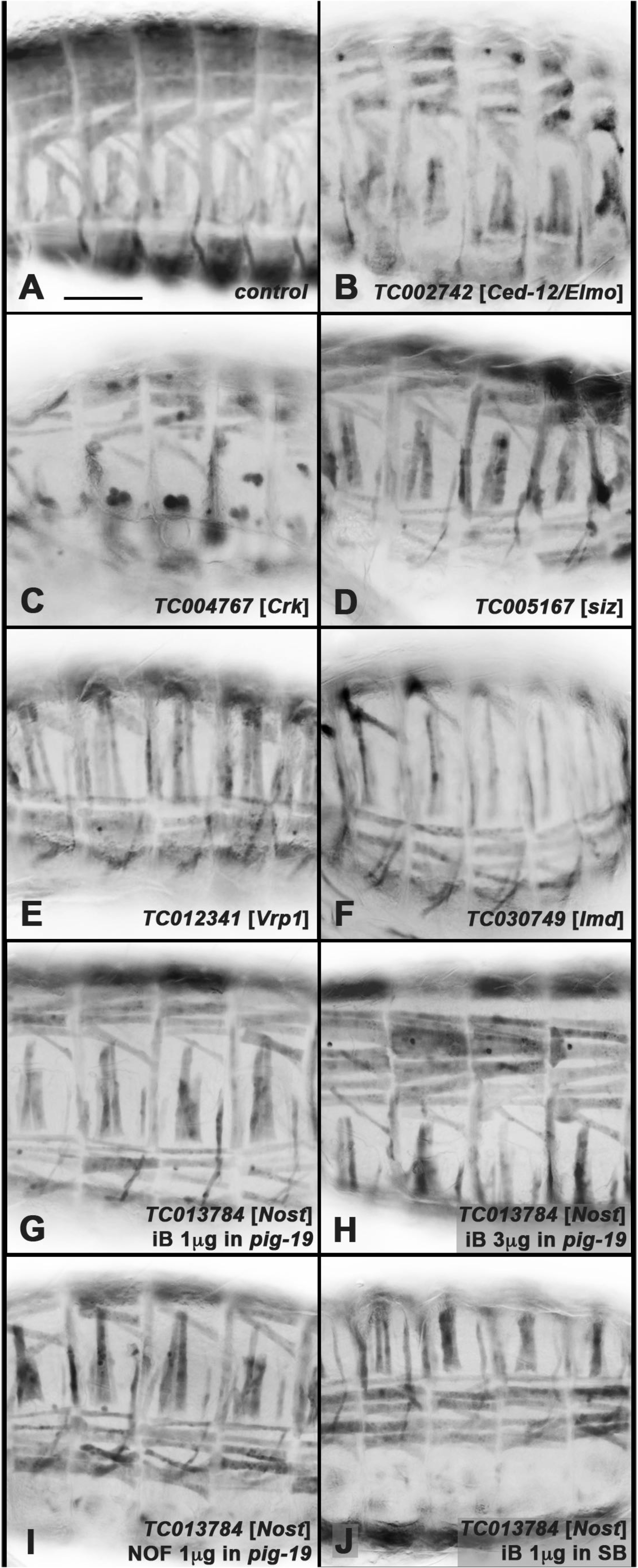
Examples of muscle phenotypes of *Tribolium* orthologs of known *Drosophila* genes required for myoblast fusion and of *TC013784* (*Tc-Nost*) Shown are lateral (A-E, G), ventral-lateral (F, I, J), or dorsal-lateral (H) views of fully developed *pig-19* embryos live imaged for EGFP. **(A)** Control embryo from uninjected female pupa. **(B)** to **(F)** Embryos from primary screen with RNAi knock-down of orthologs of known *Drosophila* genes [brackets] required for myoblast fusion as indicated. Thinner muscles and concomitantly larger distances of adjacent muscles are evident. **(G)** Embryo from primary screen (using injections of 1μg dsRNA) with RNAi knock-down of ortholog of *TC013784* (*Tc-Nost*), also exhibiting narrower muscles that are spaced apart. **(H)** Embryo from verification screen upon pupal injections of 3μg dsRNA for *Tribolium Nost* into *pig-19.* The phenotype is similar and not stronger as compared to (G). **(I)** Embryo from verification screen upon pupal injection of 1μg dsRNA for *Tribolium Nost* (non-overlapping fragment (NOF) relative to original fragment from primary screen) into *pig-19.* **(J)** Embryo from verification screen upon pupal injection of 1μg dsRNA for *Tribolium Nost* (original iBeetle fragment (iB) as in primary screen) into *SB* strain. Scale bar in A, also applicable for B – J: 100 μm.

### Knock-downs of the F-BAR domain encoding gene *Nostrin* cause similar muscle phenotypes as those of myoblast fusion genes in *Tribolium*

A new gene with a ‘thin muscle’ phenotype upon RNAi not previously implicated in myoblast fusion was *TC013784*, the ortholog of *Drosophila CG42388*, which subsequent to our screen was named after its mammalian ortholog, *Nostrin (Nost)* (Zimmermann *et al.* 2002; Zobel *et al.* 2015). The encoded Nostrin is a member of the family of F-BAR proteins that are known to regulate membrane curvature and actin turnover in a variety of contexts (Fricke *et al.* 2010; Liu *et al.* 2015; Salzer *et al.* 2017). The phenotype was present with similar strength upon injections of different amounts of the *TC013784* iB dsRNA and of a non-overlapping *Tc-Nost* dsRNA into *pig-19*, as well as upon iB dsRNA injection into the SB strain of *Tribolium castaneum* (Fig. 1G – J; c.f. Fig. 1A). In all cases, the penetrance of the phenotype was high (80 – 100% in *pig-19*, 43 – 62% in SB).

Because the observed muscle phenotype and its similarity to those of the knocked-down orthologs of the myoblast fusion genes described above were indicative of a role of *Tribolium Nost* in myoblast fusion, we tested whether *Tc-Nost (TC013784*) is expressed in the somatic mesoderm at embryonic stages when myoblast fusion is expected to occur. *In situ* hybridizations showed that *TC013784* mRNA is present at highest levels in the forming somatic muscles as well as in the CNS and posterior gut rudiment of embryos at the fully retracted germ band stage, whereas lower levels are present in epidermal cells (Fig. 2A). Hence, *TC013784* expression is compatible with a role in myoblast fusion and/or other functions in *Tribolium* muscle development. Lateral views show *TC013784* mRNA expression in subepidermal cells of the body wall and the legs, which include muscles and potentially also cells of the peripheral nervous system, as well as in specific epidermal cells (Fig. 2B).

**Figure 2.**
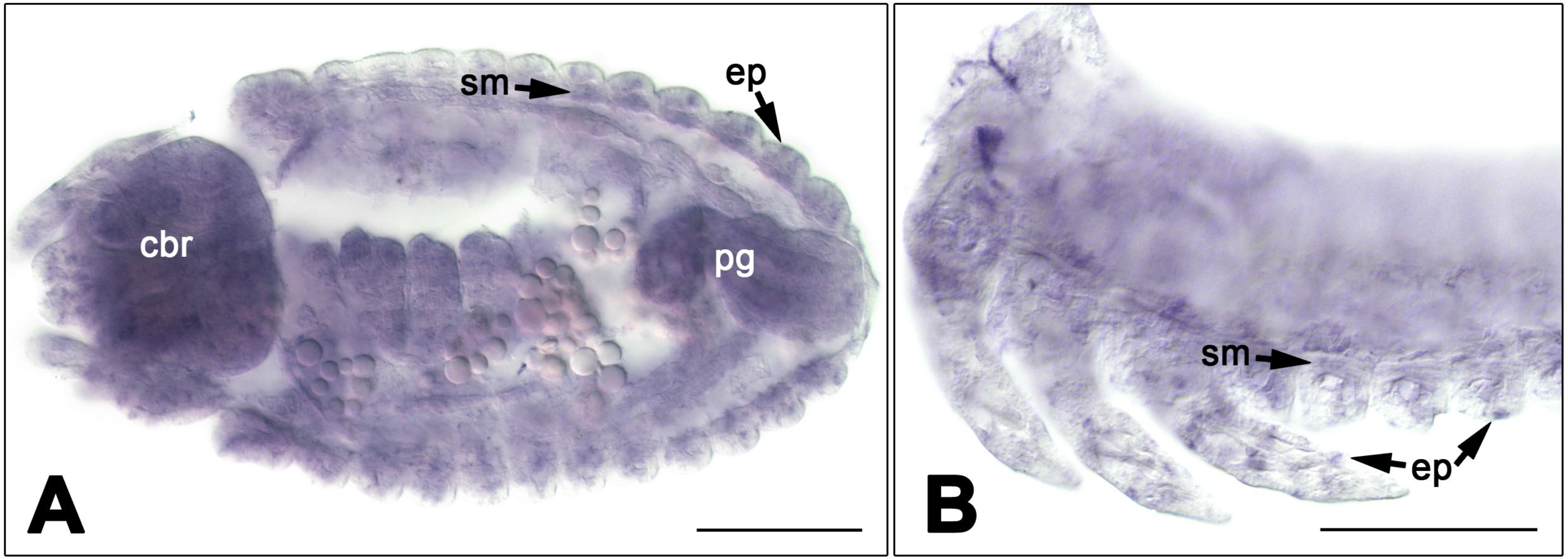
mRNA expression of *TC013784* (*Tc-Nost*) Shown are *in situ* hybridizations of embryos at the fully retracted germ band stage (A, ventral view, B, lateral view). **(A)** Higher levels of *TC013784* mRNA expression are seen in the developing somatic muscles (sm), central brain (cbr), posterior gut rudiment (pg), as well as epidermal (ep) and subepidermal cells. **(B)** *TC013784* expression is seen in somatic muscles (sm) and epidermal (ep) cells of the body wall and limbs. Scale bars: 100 μm.

### *Drosophila Nostrin* and related F-BAR domain encoding genes are expressed in the somatic and visceral mesoderm

Because of the much wider availability of immuno-histological and genetic tools in *Drosophila* we performed in-depth analyses of *Nost* and related F-BAR domain encoding genes in this insect species. As shown in Fig. 3A, *Drosophila Nost (CG42388)* mRNA is deposited maternally. The maternal transcripts vanish during stage 4 in the cellular blastoderm (except for the germ plasm and germ cells; data not shown) and zygotic expression is first seen at stage 10 in the entire mesoderm (Fig. 3B). At early stage 12, *Nost* mRNA expression becomes more restricted to segmental subsets and an anterior-posterior band of mesodermal cells, which appear to correspond to somatic and visceral mesodermal cells, respectively (Fig. 3C). To define the *Nost* mRNA expression pattern more carefully we performed fluorescent *in situ* hybridizations in conjunction with other markers for known mesodermal cell types. Double labeling for Tinman protein showed that, at stage 11, *Nost* is expressed specifically in the fusion-competent myoblasts of the trunk visceral mesoderm and the hindgut visceral mesoderm, but not in the visceral muscle founder cells and cardiogenic progenitors marked by Tinman (Fig. 3D) (Azpiazu and Frasch 1993). Double-labeling for Org-1, which marks the founder cells of visceral muscles and of a small subset of somatic muscles, confirmed the specific expression of Nost in the visceral mesodermal fusion-competent cells at stage 12, as well as in somatic mesodermal cells adjacent to the Org-1 expressing somatic muscle founder cells (Fig. 3E) (Schaub *et al.* 2012). At mid stage 12, there is a wide overlap between *Nost* mRNA and Mef2 protein expression in the somatic (and visceral) mesoderm, but not in the cardiac mesoderm (Fig. 3F) (Lilly *et al.* 1994; Nguyen *et al.* 1994). Co-stainings for *Nost* and the founder cell marker *rP298*-LacZ (aka *duf*-LacZ) indicated mutually exclusive patterns, further suggesting that *Nost* is expressed specifically in the fusion competent myoblasts of the somatic mesoderm as well (Fig. 3G, G’) (Ruiz-Gomez *et al.* 2000). This interpretation was fully confirmed by the results of *Nost in situ* hybridizations in *lameduck (lmd)* mutants, which lack fusion-competent myoblasts and do not show any *Nost* expression (Fig. 3H) (Duan *et al.* 2001). As expected for F-BAR domain proteins and shown for Nost in follicle epithelium cells (Zobel *et al.* 2015), a Nost-EGFP fusion protein localizes to the cell membranes in the somatic mesoderm (Fig. S1).

**Figure 3.**
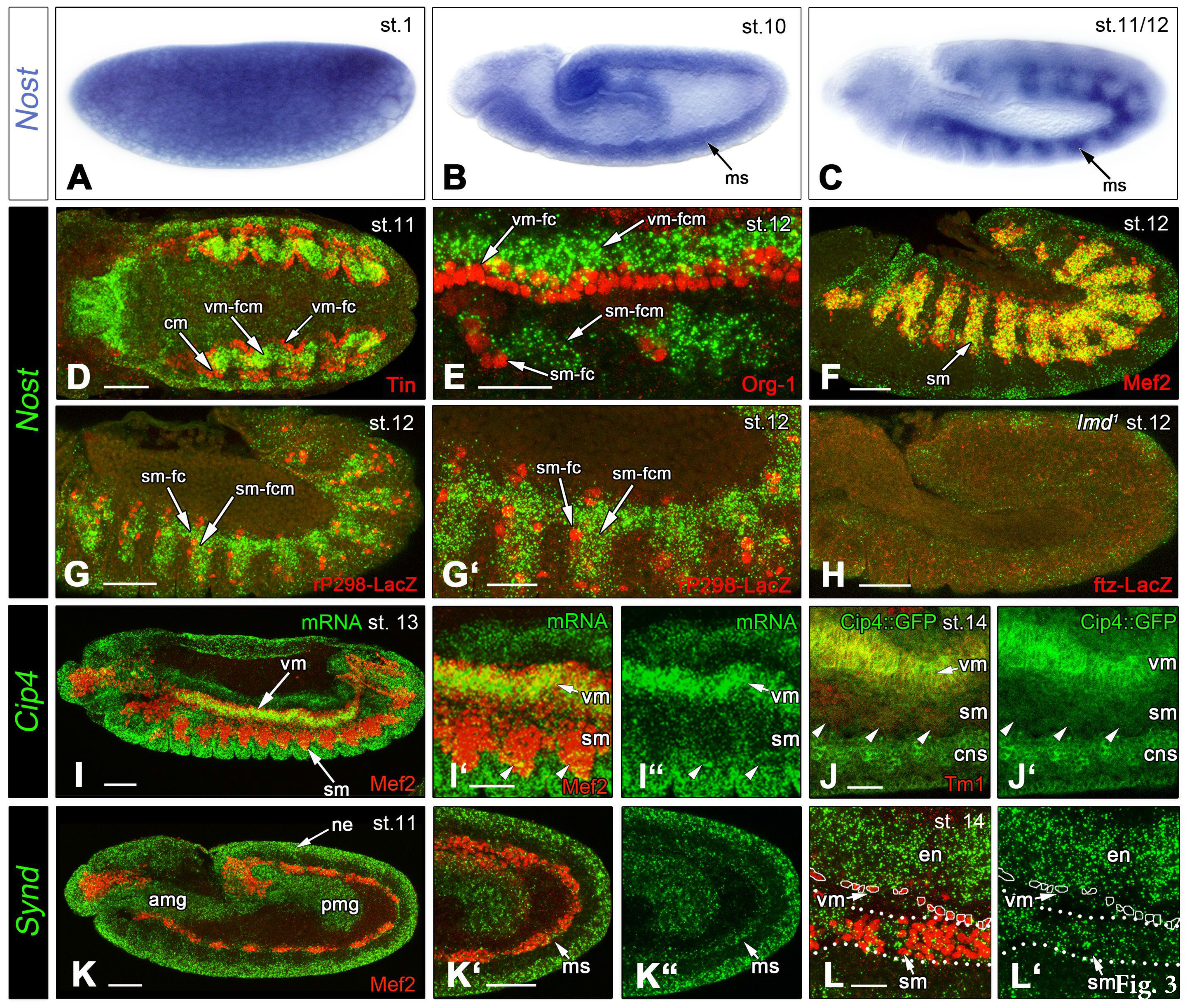
Embryonic expression of *Drosophila* F-BAR domain genes and of Cip4-YFP fusion protein. **(A)** Ubiquitous distribution of maternal *Nost* mRNA in stage 1 embryo. **(B)** Uniform zygotic mesodermal expression of *Nost* mRNA in stage 10 embryo. **(C)** Segmented mesodermal expression of *Nost* mRNA in stage 11-12 embryo. In (D) to (L’) mRNAs or fusion proteins of F-BAR domain genes are labeled in green and various tissue markers in red, as indicated. **(D)** *Nost* mRNA expression in visceral mesodermal fusion-competent myoblasts (vm-fcm) and hindgut visceral mesoderm in stage 11 embryo (dorsal view of posterior germ band). Visceral muscle founder cells (vm-fc) and cardiac mesoderm (cm) are marked by anti-Tinman (Tin) staining and lack *Nost* mRNA. **(E)** *Nost* mRNA expression in fusion-competent myoblasts of the visceral (vm-fcm) and somatic (sm-fcm) muscles (lateral high magnification view, stage 12). Visceral (vm-fc) and somatic (sm-fc) muscle founder cells are marked by anti-Org-1 staining. **(F)** *Nost* mRNA expression in fusion-competent myoblasts in the somatic mesoderm (sm; lateral view, stage 12). All somatic mesodermal cells and cardioblasts are marked by anti-Mef2 staining. **(G, G’)** *Nost* mRNA expression in fusion-competent myoblasts of the somatic muscles (sm-fcm); lateral view, stage 12 *(rP298-lacZ* line). Somatic muscle founder cells (sm-fc) are marked by rP298-LacZ enhancer trap staining (anti-βGalactosidase) and lack *Nost* mRNA. **(H)** Absent *Nost* mRNA expression in stage 12 embryo homozygous for *lmd*, which lacks sm-fcm’s. **(I)** *Cip4* mRNA expression in stage 13 embryo co-stained for Mef2 protein. **(I’, I”)** High magnification view of embryo in (I), showing *Cip4* mRNA expression in the visceral mesoderm (vm), epidermis (bottom), and more weakly in the somatic mesoderm (sm, arrow heads). **(J, J’)** High magnification view of stage 14 embryo from line tagged with YFP at native *Cip4* locus, showing Cip4-YFP expression in the visceral mesoderm (vm), central nervous system (CNS), and more weakly in the somatic mesoderm (sm, arrow heads). **(K – K”)** *Synd* mRNA expression in stage 11 embryo in mesoderm (ms), neuroectoderm (ne), and anterior as well as posterior midgut primordia (amg, pmg), with mesoderm counterstained for Mef2. **(L, L’)** *Synd* mRNA expression in stage 14 embryo (high magnification of ventral-lateral area) in somatic but not visceral mesoderm (sm, vm, counterstained against Mef2) and in endoderm (en). Scale Bars: D, F, G, H, I, K, K’, 50μm; E, G, I’, J, L, 25μm.

In addition to *Nost*, we included two other F-BAR domain encoding genes in our analysis that were characterized previously in other contexts, namely *Cip4* and *Syndapin (Synd)* (Leibfried *et al.* 2008; Fricke *et al.* 2009; Kumar *et al.* 2009b). At stages 12-13, *Cip4* mRNA is expressed prominently in the trunk visceral mesoderm, but low levels are also detected in Mef2-marked somatic mesodermal cells (in addition to ectodermal expression; Fig. 3I – I”). The same result was obtained with GFP stainings (co-stained for Tropomyosin I) of embryos from a line in which Cip4 was tagged endogenously with GFP. Similar to Nost-EGFP, Cip4-YFP fusion protein is also located at the membranes of the cells, which is most obvious for the strongly expressing visceral mesodermal, CNS, ectodermal cells, and salivary gland (Fig. 3J, J’; Fig. S1C). Low levels of Cip4-YFP are present in the somatic mesoderm. The expression of *Synd* mRNA during embryogenesis is quite broad, but co-staining with Mef2 shows that it includes the early mesodermal layer (Fig. 3K – K”) as well as the somatic mesoderm during subsequent stages (Fig. 3L, L’).

### Functionally redundant contributions of F-BAR domain genes to somatic muscle development

The specific expression of *Nost* in fusion-competent myoblasts prompted us to generate *Nost* null mutations to examine its potential functions during somatic and visceral muscle development (see Materials & Methods). In the two alleles obtained, all (*Nost*^ff016^) or almost all (*Nost*^df004^) protein-coding exons of each isoform were deleted (Fig. S2). Homozygous flies for both alleles were fully viable, fertile, and lacked any obvious defects, including in locomotion, as was also shown with a *Nost* allele presumably identical to *Nost*^df004^ that was made in parallel (Zobel *et al.* 2015). Furthermore, embryos collected from homozygous *Nost* mutant strains, which therefore lacked both the maternal and the zygotic activity of *Nost*, did not exhibit any defects in their somatic muscle patterns (Fig. 4D, cf. Fig. 4A, B). Likewise, embryos from crosses of homozygous null mutant flies for *Cip4*, which completely lack *Cip4* activity (Fricke *et al.* 2009), also did not show any somatic muscle phenotype (Fig. 4E), and neither did homozygous *Synd* null mutant embryos (which do contain maternal products, as *Synd* is homozygously larval lethal (Kumar *et al.* 2009a) and fertile homozygous *Synd*^Δex22^ mutant females could not be obtained) (Fig. 4F).

**Figure 4.**
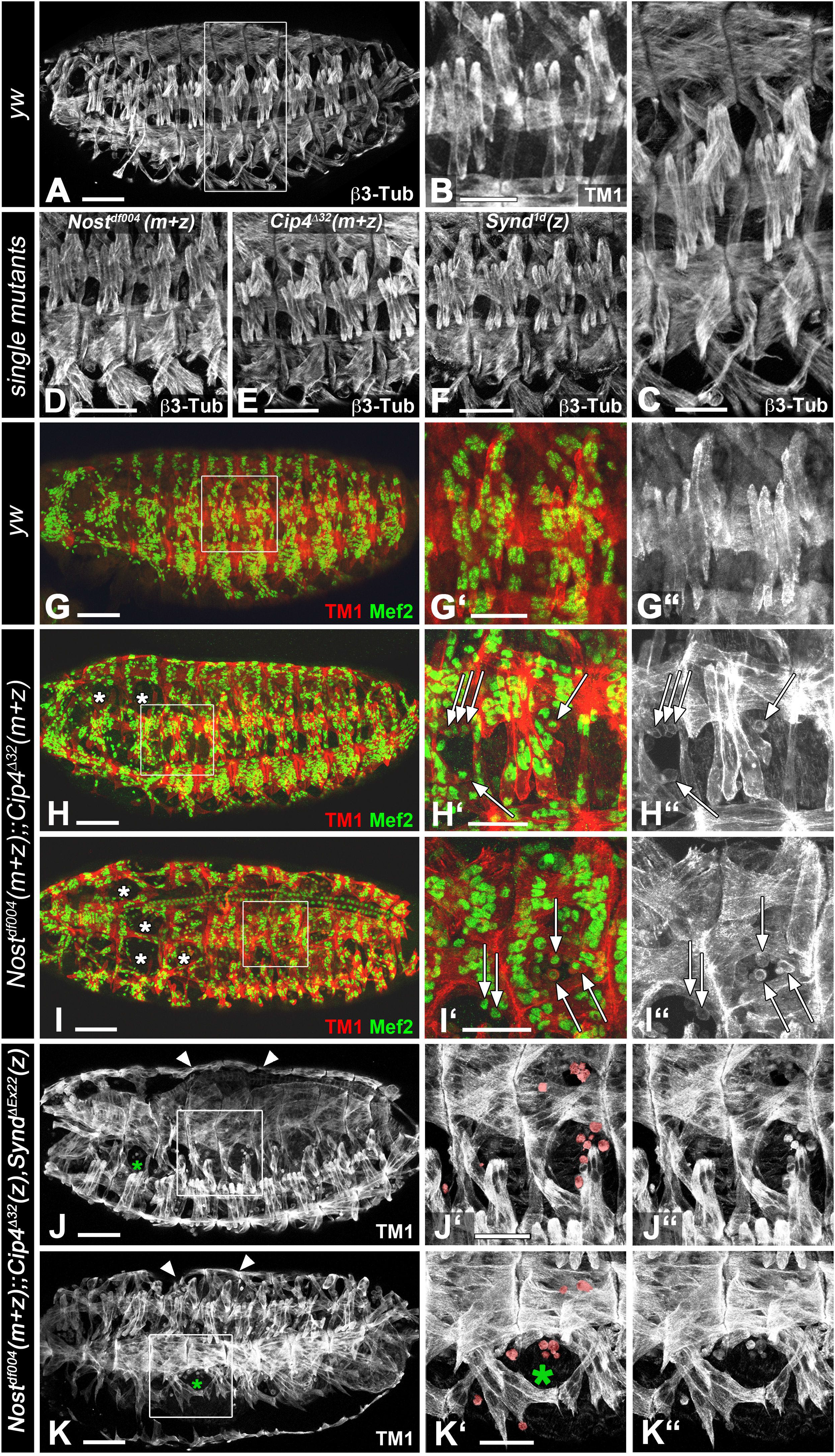
Embryonic somatic muscle phenotypes in mutants for F-BAR domain genes. Shown are stage 16 embryos derived from *inter se* crosses of homozygous single mutant parents for *Nost* (D) or *Cip4* (E), homozygous mutant embryos from heterozygous *Synd* mutant parents (F) and from homozygous *Nost Cip4* double mutant escaper parents (H – I”), and embryos homozygous for mutations in all three genes from homozygous *Nost* mutant parents that were heterozygous for *Cip4* and *Synd* (J – K”). (**A**) Somatic muscle pattern in control embryo (yw) stained for β3-tubulin. **(B)** High magnification view of lateral muscles in two abdominal segments of control embryo stained for tropomyosin I (TM1). **(C)** High magnification view of two abdominal segments from embryo in A (boxed). **(D, E, F)** Views of normal ventral and lateral muscles of four abdominal segments from maternally + zygotically mutant (m+z) *Nost*^df004^, *Cip4^A32^*, and of zygotic *Synd^1d^* mutant, respectively, all stained for β3-tubulin. **(G – G”)** Somatic muscle pattern in control embryo (yw) stained for tropomyosin I (TM1, red) and Mef2 (green) (G’, high magnification view of boxed area in G; G” single channel for TM1). **(H – I”)** Somatic muscle pattern in maternally + zygotically (m+z) *Nost*^df004^;;*Cip4*^D32^ double mutants stained and depicted as in (G – G”). Arrows indicate mononucleated myoblasts. **(J – K”)** Somatic muscle pattern in maternal + zygotic (m+z) *Nost*^df004^;;zygotic (z) *Cip4^A32^ Synd*^Δex22^ triple mutants stained for TM1 and depicted as in (G – G”). Arrows indicate mononucleated myoblasts (highlighted in J’, K’), asterisks an example of area with missing muscles, and arrow heads a dorsal bulge due to expanded midgut. Abbreviations: m+z, zygotically homozygous and lack of maternal contribution; z, zygotically homozygous with presence of maternal contribution. Scale Bars: A, D, E, F, G, H, I, J, K, 50μm; B, C, G’, H’, I’, J’, K’, 25μm.

Because of the possibility of functional redundancies among these different F-BAR proteins we examined *Nost Cip4* double mutants and *Nost Cip4 Synd* triple mutants at stage 16 for embryonic muscle phenotypes. Synergistic activities of *Nost* and *Cip4* have already been demonstrated by the appearance of egg chamber defects in *Nost Cip4* double mutants (Zobel *et al.* 2015). In addition, simultaneous knockdowns of *Nost* and *Cip4* led to increased duplicated and frequent multiple wing hair phenotypes as compared to *Nost* mutants that lack any wing hair phenotype and *Cip4* mutants that show duplicated wing hairs at lower frequency (Zobel *et al.* 2015). As shown in Fig. 4H – I” (c.f. Fig. 4C, G – G”), mutant embryos completely lacking both *Nost* and *Cip4* products (see Materials & Methods) indeed displayed frequent muscle defects. In several segments, certain muscle fibers were missing or strongly reduced in size, and instead, mononucleated myoblasts were present at the corresponding positions. These can be detected as Tropomyosin I-positive cells that contain a single Mef2-positive nucleus each (Fig. 4H’, H”, I’, I”; c.f. Fig. 4C, G’, G”). Whereas control embryos do not contain any unfused myoblasts at stage 16, unfused myoblasts were present inappropriately in all *Nost* (m+z) *Cip4* (m+z) double mutant embryos at this stage, and about half of these had muscles missing in one to four segments (Table S1A). We did not detect any preferential distribution of these muscle defects with respect to specific segmental or muscle identities. No defects were visible in the dorsal vessel (Fig. 4I).

We found that, similar to *Cip4* and *Nost, Cip4* and *Synd* also have synergistic activities during epithelial planar polarity and wing hair formation (Fig. S3). Therefore we examined the embryonic musculature in *Cip4 Synd* double mutant embryos lacking the zygotic activities of both *Cip4* and *Synd.* Although no muscle defects were detected in these embryos (Fig. S3), such defects were present in embryos in which the zygotic activities of *Synd* and *Cip4* were missing together with both the maternal and zygotic activities of *Nost* (Fig. 4J – K”; c.f. Fig. 4C, G”). Like in *Nost* (m+z) *Cip4* (m+z) double mutant embryos, muscle fibers were variably missing and, unlike in the controls, unfused myoblasts were seen in late stage embryos. In the triple mutants *(Nost* (m+z) *Cip4* (z) *Synd* (z)) this phenotype appeared to be slightly more severe, even though *Cip4* activity is only removed zygotically in these embryos. This indicates that *Synd* also contributes to normal somatic muscle development, in cooperation with *Nost* and *Cip4.* Altogether, the observed phenotypes suggested a role of these F-BAR proteins in the process of myoblast fusion. To describe the muscle phenotypes more quantitatively in the different genetic backgrounds, we counted syncytial nuclei at consecutive developmental stages of double and triple mutant embryos. For this analysis we used the well-characterized dorsal muscle DA1 (aka, M1) that expresses Even-skipped, which was used in combination with lamin as a marker for the nuclei. In control embryos of stages 14, early 15, late 15, and 16, the increasing numbers of nuclei counted within the DA1 syncytia closely matched the numbers determined previously by Bataille *et al.* (2010). By contrast, in both double and triple mutant embryos, already at stage 14 the numbers of nuclei within the DA1 syncytia were lower as compared to the controls (Fig. 5, Table S1B), suggesting that about one less myoblast fusion event had occurred in these mutants at this stage. These differences increased further until late stage 15 and stage 16, when the double and triple mutant embryos on average contained about two to three fewer nuclei within the DA1 muscle fibers. These data confirm that *Nost* and *Cip4* have cooperative activities in promoting myoblast fusion. As the difference between the double and triple mutants were not statistically significant, it is not clear whether *Synd* indeed contributes to the functions of *Nost* and *Cip4* during this process.

**Figure 5.**
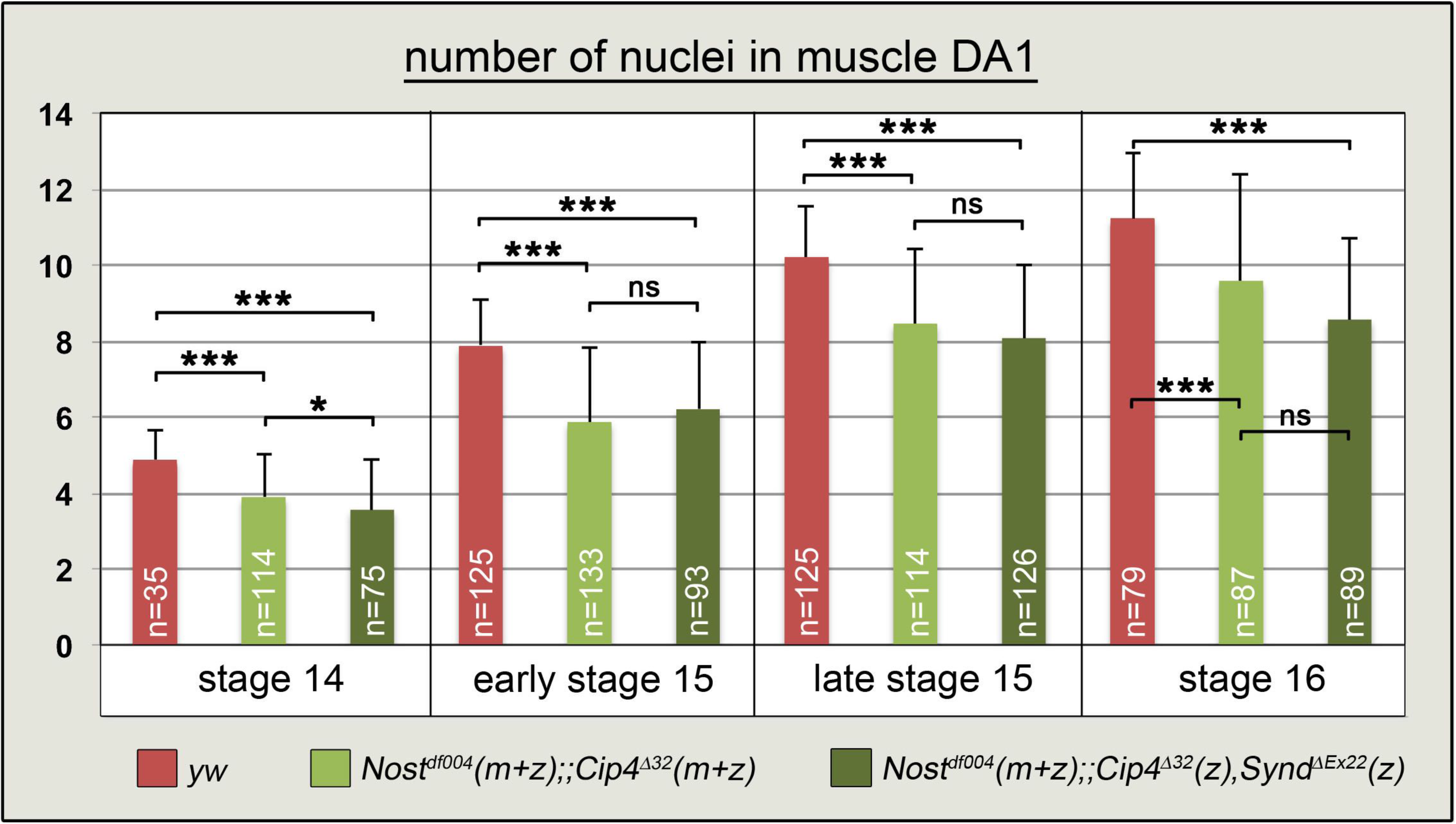
Quantification of nuclei within muscle DA1 syncytia of control, *Nost Cip4* double, and *Nost Cip4 Synd* triple mutant embryos. Even-skipped + Lamin stained nuclei of muscle DA1 syncytia, counter stained for tropomyosin I, were counted at stages 14, early and late stage 15, and stage 16 in *yw* control embryos and in embryos of the genotypes *Nost*^df004^(m+z);;*Cip4*^D32^ (m+z) and *Nost*^df004^(m+z);;*Cip4*^Δ32^(m+z),*Synd*^Δex22^(z) (***, p <0,0005; *, p <0.05; ns, differences not significant; m, maternal; z, zygotic).

### F-BAR domain encoding genes are required for normal embryonic midguts and for midgut muscle morphogenesis during metamorphosis

In embryos lacking the zygotic activities of both *Synd* and *Cip4*, and likewise in embryos lacking zygotic *Synd* together with zygotic *Cip4* and maternal + zygotic *Nost*, a subtle but consistent phenotype was seen in the midgut. In embryos with these genetic backgrounds, the anterior chamber was slightly expanded, thereby causing a bulge at the dorsal side of the embryos (Fig. 6 B, c.f. Fig.6A; Fig. S3E; c.f. Fig. S3D; Fig. 4J, K; c.f. Fig. 4G). This phenotype was not present in embryos that were only homozygous for *Synd*, thus indicating that *Cip4* and potentially *Nost* cooperate with *Synd* during normal midgut morphogenesis. Because disruptions in midgut morphology are often due to developmental defects in the visceral musculature (Lee *et al.* 2005) and because particularly *Nost* and *Cip4* showed prominent expression in the visceral mesoderm, we examined whether loss of these F-BAR domain proteins or of Synd caused any gut muscle phenotypes.

**Figure 6.**
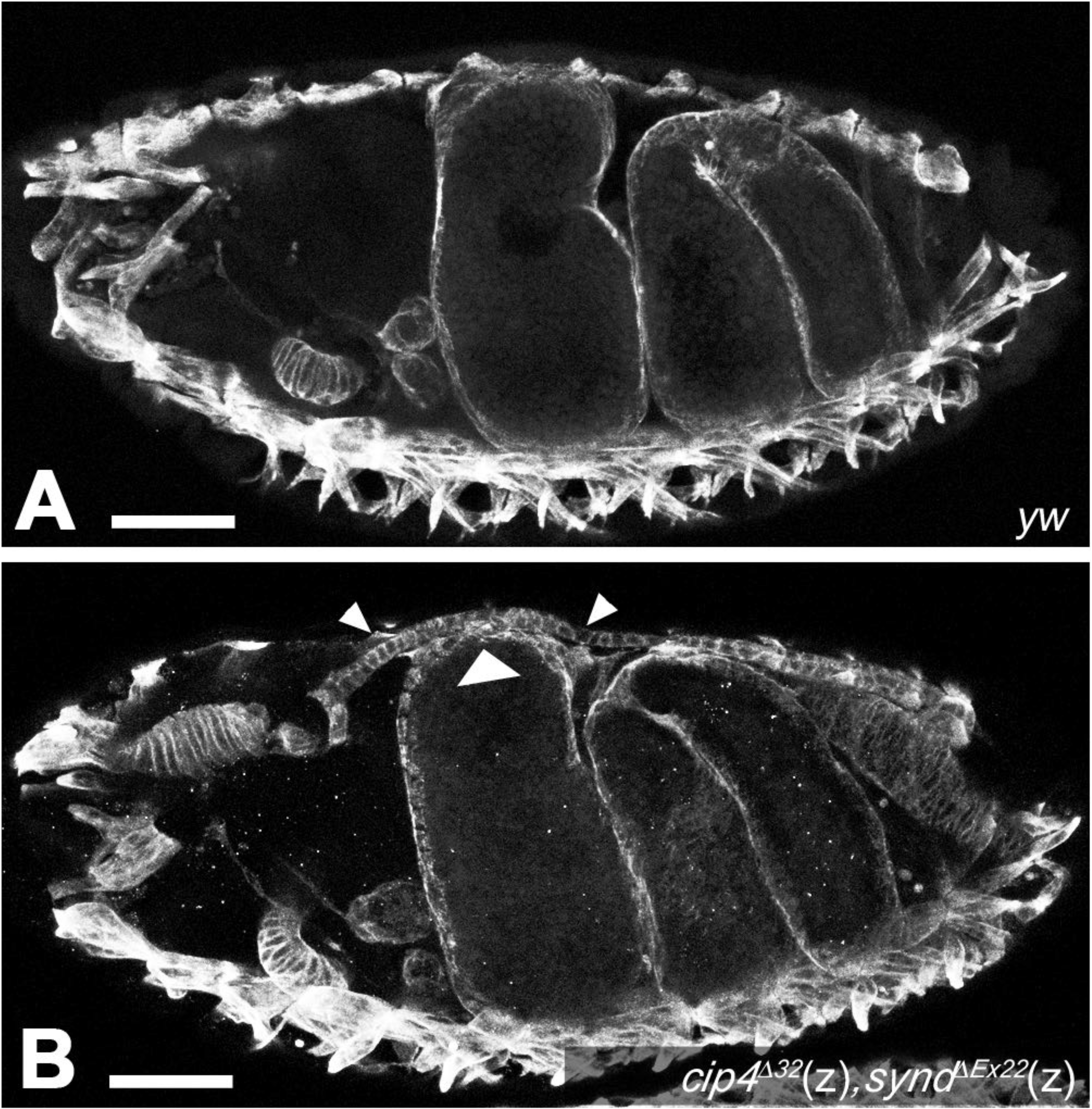
Midgut morphology defects in *Cip4 Synd* double mutant embryos. **(A)** Optical section of stage 16 *yw* control embryo (anti-Tropomyosin I). **(B)** Optical section of stage 16 homozygous *cip4*^Δ32^(z), *synd*^ΔEx22^(z) embryo. The anterior midgut chamber is slightly expanded, thus creating a bulge at the dorsal side of the embryo (arrow heads). Abbreviation: z, zygotically mutant (homozygous). Scale Bars: 50μm.

In embryos (and, likewise, in larvae), none of the mutant backgrounds described above displayed any overt gut muscle phenotypes, which is in line with the relatively mild alterations in their midgut morphologies. By contrast, strong defects were seen in midguts of adults with some of these mutant backgrounds. Normal adult midguts are ensheathed by an orthogonal network of binucleated circular and multinucleated longitudinal visceral muscles, which run in parallel and are arranged equidistantly to their neighboring fibers (Strasburger 1932; Klapper 2000). The same pattern as in control flies is observed in midguts of *Nost* (m+z) mutant flies (Fig. 7B, c.f. Fig. 7A). However, in *Cip4* (m+z) mutant flies, particularly the longitudinal muscle fibers show strongly disrupted morphologies. As shown in Fig. 7C – C”, in the absence of Cip4 the longitudinal muscle fibers display numerous branches, particularly near their ends, which often contact neighboring longitudinal fibers. In addition, the longitudinal fibers appear shorter, as unlike in the wild type they do not span large extents of the length of the midgut, and they are also not arranged strictly in parallel. In *Nost* (m+z) *Cip4* (m+z) double mutant flies (see Materials & Methods), analogous but even more severe disruptions of midgut muscle morphologies are observed. In these flies, the ends of the longitudinal muscle fibers are even more frayed and some fibers are split towards their middle portions (Fig. 7D – D”). Furthermore, the arrangement and thickness of the longitudinal fibers is irregular. In *Cip4* (m+z) mutants and, even more often, in *Nost* (m+z) *Cip4* (m+z) double mutants the normally parallel arrangement of the circular midgut muscles is also disrupted (Fig. 7C – C”, Fig, 7D – D”). Their distances vary widely and often fibers cross over each other. Closer inspection showed that the abnormal feathered extensions of the longitudinal fibers tend to contact the circular fibers at their ends, and it appears that the circular fibers are being pulled in anterior-posterior directions by the contractions of the attached longitudinal fibers (Fig. 6C’ – D”). Hence, the circular muscle phenotype may be largely secondary to the observed longitudinal muscle phenotype. These data show that *Cip4* and *Nost* cooperate during the process of longitudinal midgut muscle metamorphosis. A contribution of *Synd* cannot readily be tested because homozygous *Synd* mutant flies are not viable.

**Figure 7.**
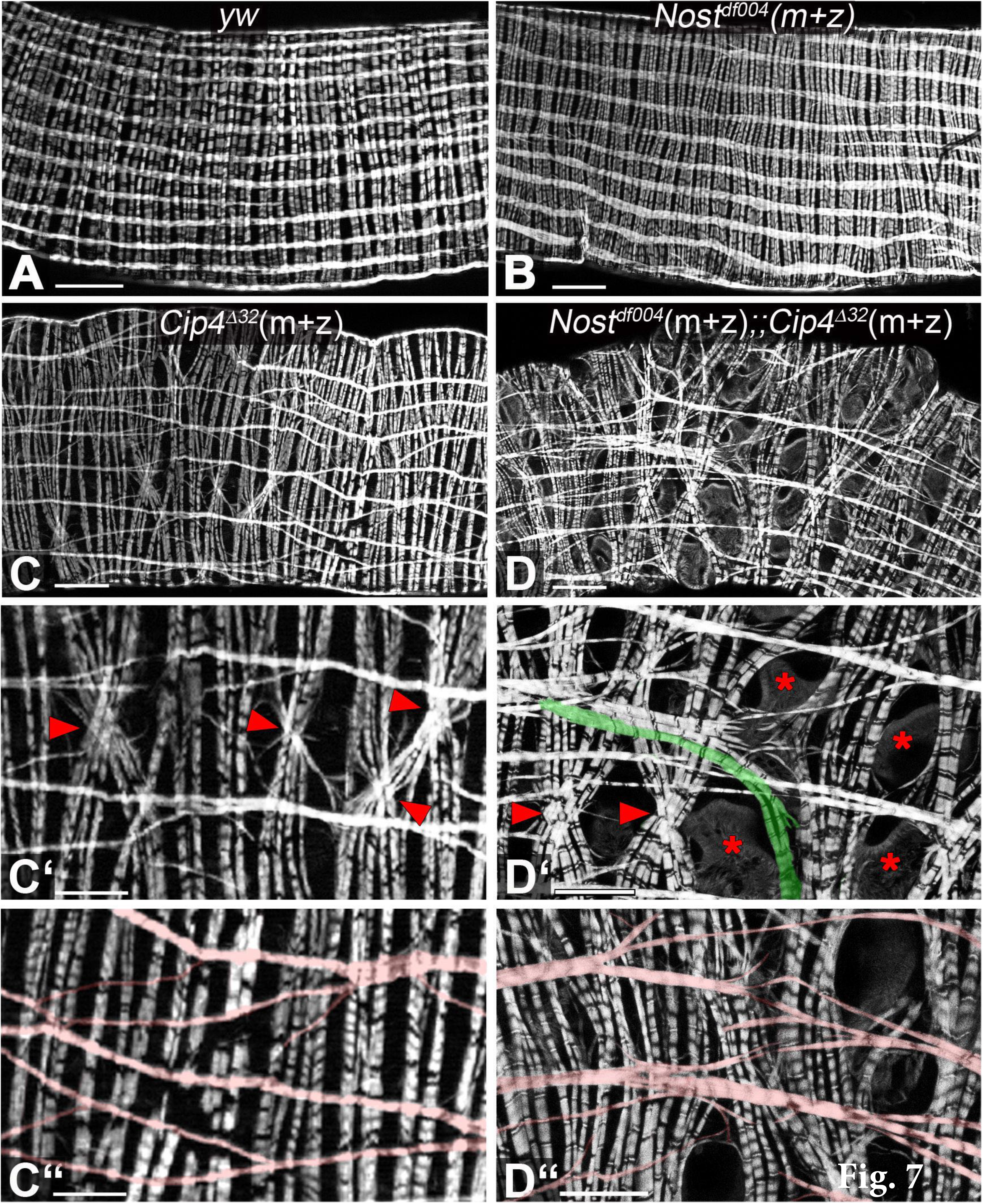
Midgut muscle phenotypes in adult escaper flies singly mutant for *Nost* or *Cip4* and doubly mutant for *Nost Cip4*. Shown are portions of midguts stained with phalloidin from young adults. **(A)** Orthogonal longitudinal and circular muscles of*yw* control fly. **(B)** Normal longitudinal and circular muscle pattern in midgut from *Nost*^df004^(m+z) fly. **(C)** Disrupted longitudinal and circular muscle pattern in midgut from *Cip4*^Δ32^(m+z) fly. **(C’, C”)** High magnification views of midgut in (C). Crossed circular muscles are indicated by arrow heads in (C’) and aberrantly split and mis-oriented longitudinal muscles are highlighted pink in (C”). **(D)** Disrupted longitudinal and circular muscle pattern in midgut from *Nost*^df004^(m+z);;*Cip4*^Δ32^(m+z) fly. **(D’)** High magnification view of midgut in (D). Crossed circular muscles are indicated by arrow heads, large gaps by asterisks, and longitudinal muscle aberrantly curved towards circular muscles is highlighted in green. **(D”)** High magnification view of different midgut with genotype as in (D). Aberrantly split and mis-oriented longitudinal muscles are highlighted in red. Abbreviations: m+z, zygotically homozygous and lack of maternal contribution (see legend fig. 4). Scale Bars: A – D, 50μm; C’ – D”, 25μm.

## Discussion

### F-BAR proteins promote myoblast fusion

Prompted by the observed thin-muscle phenotype upon *Tc-Nostrin (TC013784)* knockdown and the highly specific expression of *Dm-Nostrin* in fusion-competent myoblasts of embryonic somatic and visceral muscles, we focused on the characterization of the potential roles of Dm-Nostrin (Nost) and two structurally related proteins, Cip4 and Syndapin (Synd), in *Drosophila* muscle development. All three proteins contain F-BAR domains within their N-terminal half and an SH3 domain at their C-terminus (Fricke *et al.* 2009; Kumar *et al.* 2009a; Zobel *et al.* 2015). These F-BAR proteins belong to the NOSTRIN, CIP4, and PACSIN subfamilies of the F-BAR protein superfamily and are the only representatives of this subfamily in *Drosophila*, and likewise, in *Tribolium* (although there are two *Tc-Cip4* paralogs, see Materials & Methods). Members of this F-Bar protein subfamily are known to provide a molecular link between the plasma membrane or nascent vesicular membranes and actin dynamics (Liu *et al.* 2015; Salzer *et al.* 2017). Chiefly, the interaction of the homodimeric crescent-shaped F-BAR domains of these proteins with membrane phospholipids creates membrane curvatures, and their SH3 domains interact with the Wiskott-Aldrich syndrome protein (WASp), neural (N)-WASp, WASP family verproline-homologous protein (WAVE), and dynamin. The ensuing activation of these proteins leads to the binding of actin-related protein 2/3 (Arp2/3) which, in turn, promotes actin nucleation and polymerization. Actin polymerization then can further propel membrane curvatures, which in the case of the NOSTRIN, CIP4, and PACSIN family proteins has been reported to promote the formation of filopodia, lamellopodia, podosomes, invadopodia, and to stimulate endocytosis (Chen *et al.* 2012; Liu *et al.* 2015; Salzer *et al.* 2017). Cellular events with these characteristics are also known to be hallmarks during *Drosophila* myoblast fusion, particularly in fusion-competent myoblasts, in which *Dm-Nostrin* is expressed (Önel and Renkawitz-Pohl 2009; Kim *et al.* 2015; Deng *et al.* 2017). Thus, during the earliest steps of myoblast fusion, fusion-competent myoblasts extend filopodia to the muscle founder cells before attaching to them (Ruiz-Gomez *et al.* 2000). The actual fusion process is driven to a large part by the formation of a dense F-actin focus surrounded by a fusion-restricted myogenic-adhesive structure (FuRMAS) in fusion-competent myoblasts, which propels an invadopodia-like membrane protrusion into the attached founder myoblast or nascent myotube (Önel and Renkawitz-Pohl 2009; Kim *et al.* 2015; Deng *et al.* 2017). This process is thought to provide the key force for membrane rupture and cell fusion (Sens *et al.* 2010). F-actin polymerization is activated downstream of the activated Ig domain receptors Sticks-and stones (Sns) and Hibris (Hbs) upon their engagement with the extracellular domains of the related receptors Kirre (aka Dumbfounded, Duf) and Roughest (Rst) on the surface of the founder myoblast. Links between the intracellular domains of active Sns/Hbs are provided by the adaptor proteins Dock and Crk, which bind to activated Sns and Hbs and through their SH3 domains interact with WASp and the WASp regulator Verprolin 1 (Vpr1; aka Solitary, Sltr). In turn, these nucleate linear and branched actin polymerization via activation of Arp2/3, which is additionally regulated by activated WAVE (aka SCAR), the WASp family member WHAMY, and by the formin Diaphanous (Dia) (Önel and Renkawitz-Pohl 2009; Kim *et al.* 2015; Brinkmann *et al.* 2016; Deng *et al.* 2017). As our data demonstrate a contribution of F-BAR proteins, particularly Nostrin and Cip4, to the process of myoblast fusion, it is conceivable that these proteins provide an additional, perhaps receptor-independent link between the plasma membrane and actin polymerization at the fusion site within fusion-competent myoblasts. In addition to their activation of actin polymerization, they could influence membrane bending at the fusion site directly by binding to the plasma membrane, and help coordinating membrane bending and the formation of the F-actin focus. However, attenuation of fusion efficiency even after the complete elimination of both Nost and Cip4 (and additionally of zygotic Synd) is relatively mild as compared to the generally stronger block of myoblast fusion in mutants for the various actin nucleators and their upstream activators (unless there are strong maternal contributions)(e.g., Richardson *et al.* 2007; Gildor *et al.* 2009; Kaipa *et al.* 2013). This indicates that these F-BAR proteins play a supporting rather than an essential role during this process. In addition, as shown herein, there is functional redundancy among different F-BAR proteins during this process. Overall, this and published information support the view that the system has a significant amount of back-up pathways built in. In *Tribolium*, we have not attempted any double or triple knock-downs of the different subfamily members to determine whether this would cause more severe muscle phenotypes.

As F-BAR proteins including Nostrin, Cip4, and Syndapin in both mammals and *Drosophila* have been shown to regulate receptor-mediated endocytosis (Kessels and Qualmann 2002; Icking *et al.* 2005; Itoh *et al.* 2005; Leibfried *et al.* 2008; Fricke *et al.* 2009; Feng *et al.* 2010; Zobel *et al.* 2015; Sherlekar and Rikhy 2016), this mode of action could also be involved in promoting myoblast fusion. In *Drosophila* myoblast fusion, there is evidence that local clearance of N-cadherin at the fusion site by endocytosis in fusion-competent cells and nascent myotubes is needed prior to fusion to allow progression of the fusion process (Dottermusch-Heidel *et al.* 2012). Perhaps related to this observation, *Nost* and *Cip4* were shown to cooperate in sequential steps of endocytotic E-Cadherin membrane turnover in the *Drosophila* thoracic epithelium and in developing egg chambers, which in the latter case is important for proper germline cell adhesion, egg chamber encapsulation by follicle cells, and normal fertility (Zobel *et al.* 2015). In growing myotubes, endocytosis appears to be involved also in the clearance of Sns (but not Duf) in addition to N-cadherin, which may be beneficial for efficient later rounds of myoblast fusion (Haralalka *et al.* 2014). In future experiments, these processes could be monitored in *Nost Cip4* double or *Nost Cip4 Synd* triple mutant embryos to determine a possible role of these F-BAR proteins in endocytotic events during myoblast fusion.

In mouse, several observations from *in vitro* models have pointed to the involvement of F-BAR and other BAR superfamily proteins in myoblast fusion. (George *et al.* 2014) reported that the CIP4 subfamily member Toca-1 is required for normal myoblast fusion and myotube formation in differentiating C2C12 cells, which appears to involve downstream activation of the actin regulator N-WASp. In an *in vitro* model for cell-to-cell fusion initiated by protein fusogens of influenza virus and baculovirus, curvature generating proteins, including the F-BAR domain protein FCHo2 as well as GRAF1 that contains a C-terminal SH3 domain in addition to the N-terminal Bar domain and central RhoGAP and pleckstrin homology (PH) domains, were shown to promote syncytium formation (Richard *et al.* 2011). GRAF1 is enriched in skeletal muscle and was reported to promote terminal differentiation and myoblast fusion of C2C12 cells, which involves its Rho-GTPase activating function for actin remodeling and BAR domain-dependent membrane sculpting (Doherty *et al.* 2011). Myoblasts isolated from GRAF1 knock-out mice and regenerating muscles in GRAF1^-/-^ mice showed reduced myoblast fusion (Lenhart *et al.* 2014). In addition to its influence on the actin metabolism, this function may be mediated by regulating vesicular trafficking of the fusogenic ferlin proteins to promote membrane coalescence (Lenhart *et al.* 2014). *Drosophila* Graf is known to regulate hematopoiesis through endocytosis of EGFR, but its potential expression in the somatic mesoderm and any role in myoblast fusion have not been examined (Kim *et al.* 2017). Yet another Bar family member, the N-Bar domain protein Bin3, has also been implicated in mouse myoblast fusion, as myoblasts from Bin3^-/-^ mice show a reduced fusion index and Bin KO mice show delayed regeneration upon injury (Simionescu-Bankston *et al.* 2013).

### F-BAR proteins are important during adult visceral muscle morphogenesis

The strongest muscle phenotypes of *Drosophila* F-BAR gene mutants are present in the longitudinal midgut muscles of adult flies, which instead of being linear and arranged in parallel are highly branched at their ends, connected to their neighbors, and oriented irregularly. In this case, the phenotype is already seen in *Cip4* single mutants but is enhanced in *Nost Cip4* double mutants, similar to the situation in wing hair formation (Zobel *et al.* 2015). The phenotype is most likely explained by the drastic events of cellular remodeling of the longitudinal midgut muscles during metamorphosis. During pupariation, the larval longitudinal muscles dedifferentiate, first forming numerous cytoplasmic projections that are shed (Klapper 2000), then fragmenting into smaller syncytia and finally into mononucleated myoblasts (Aghajanian *et al.* 2016), DS & MF, unpublished data). At this stage of maximal dedifferentiation the myoblasts are connected to each other by a network of fine filopodia. The reconstitution of the visceral muscle syncytia is accompanied by a progressive disappearance of the lateral extensions to neighboring cells and syncytia (DS & MF, unpublished data). Ultimately parallel, multinucleated fibers are re-established that very much resemble the original larval longitudinal midgut muscles. Because the longitudinal midgut muscles in *Cip4* and even more so in *Nost Cip4* mutant flies are shorter, display frayed ends connected to neighboring muscles, and are not neatly arranged in parallel, we propose that these F-BAR domain proteins act especially during the steps of redifferentiation. Apart from myoblast fusion, we propose that shape changes and removal of the extensive filopodial connections during these extreme remodeling events involve coordinated interactions between membranes and actin turnover, as well as regulated endocytosis, in which these F-BAR proteins are likely involved. Future experiments with fluorescent plasma membrane reporters in longitudinal muscles in *Nost Cip4* mutant pupae can test these possibilities, although so far our attempts in this direction were hampered by the very low fertility of these double mutants, which has limited the number of developmental stages that could be examined.

### Conclusion

Aiming to utilize the identified *Tribolium* genes from the iBeetle screen for gaining new insight into *Drosophila* muscle development, we demonstrated that F-BAR domain proteins, particularly Nostrin and Cip4, play roles in myoblast fusion during embryonic somatic muscle development and during visceral muscle remodeling at metamorphosis. The examination of orthologs of additional genes with muscle phenotypes identified in the iBeetle screen will likely further advance our understanding of muscle development in *Drosophila* and other species.

## Acknowledgments

We acknowledge support by the German Research Foundation (DFG) for the *iBeetle* project (FOR1234). We thank Sven Bogdan for discussions and the exchange of at the time unpublished materials. We also thank the Bloomington Stock Center, the Kyoto Stock Center, and the CDRC Stock Center for *Drosophila* stocks, the *Drosophila* Genomics Resource Center for clones, and Renate Renkawitz-Pohl, Hanh Nguyen, and the Developmental Studies Hybridoma Bank (DHSB) for antibodies.

## Supplementary files

**Figure S1.**
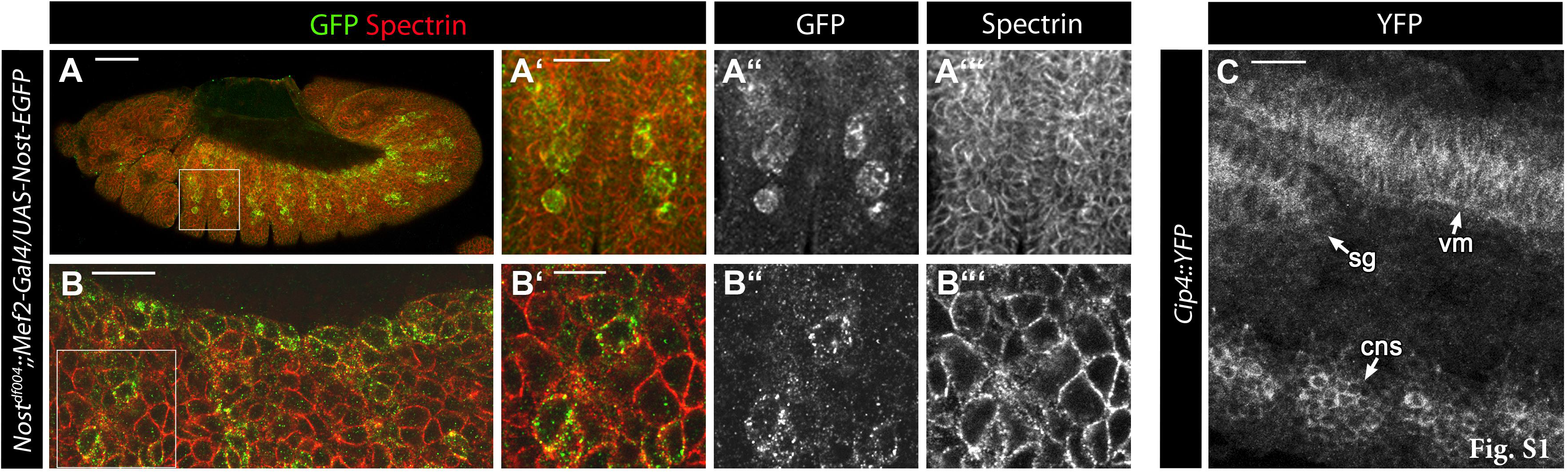
Localization of Nost-EGFP and Cip4-YFP fusion proteins. (**A – A”’)** and (**B – B”’**) Homozygous *Nost*^df004^ null mutant embryos with *UAS-Nost-EGFP* driven by *Mef2-GAL4* in the somatic mesoderm and stained with anti-GFP and anti-Spectrin antibodies. High magnification views in (A’ – A”) and (B’ – B”) of boxed areas from (A) and (B), respectively, of somatic mesodermal layers show predominant colocalization of the two proteins within myoblasts, thus indicating localization of Nost-EGFP underneath the plasma membrane. **(C)** High magnification view of stage 14 embryo from line tagged with YFP at native *Cip4* locus, showing Cip4-YFP localization around the circumference of cells in the visceral mesoderm (vm), central nervous system (CNS), and salivary gland (sg). Scale bars: A, 50μm; A’ – A””, B, C, 20 μm; 20μm; B’ – B’”, 10 μm.

**Figure S2.**
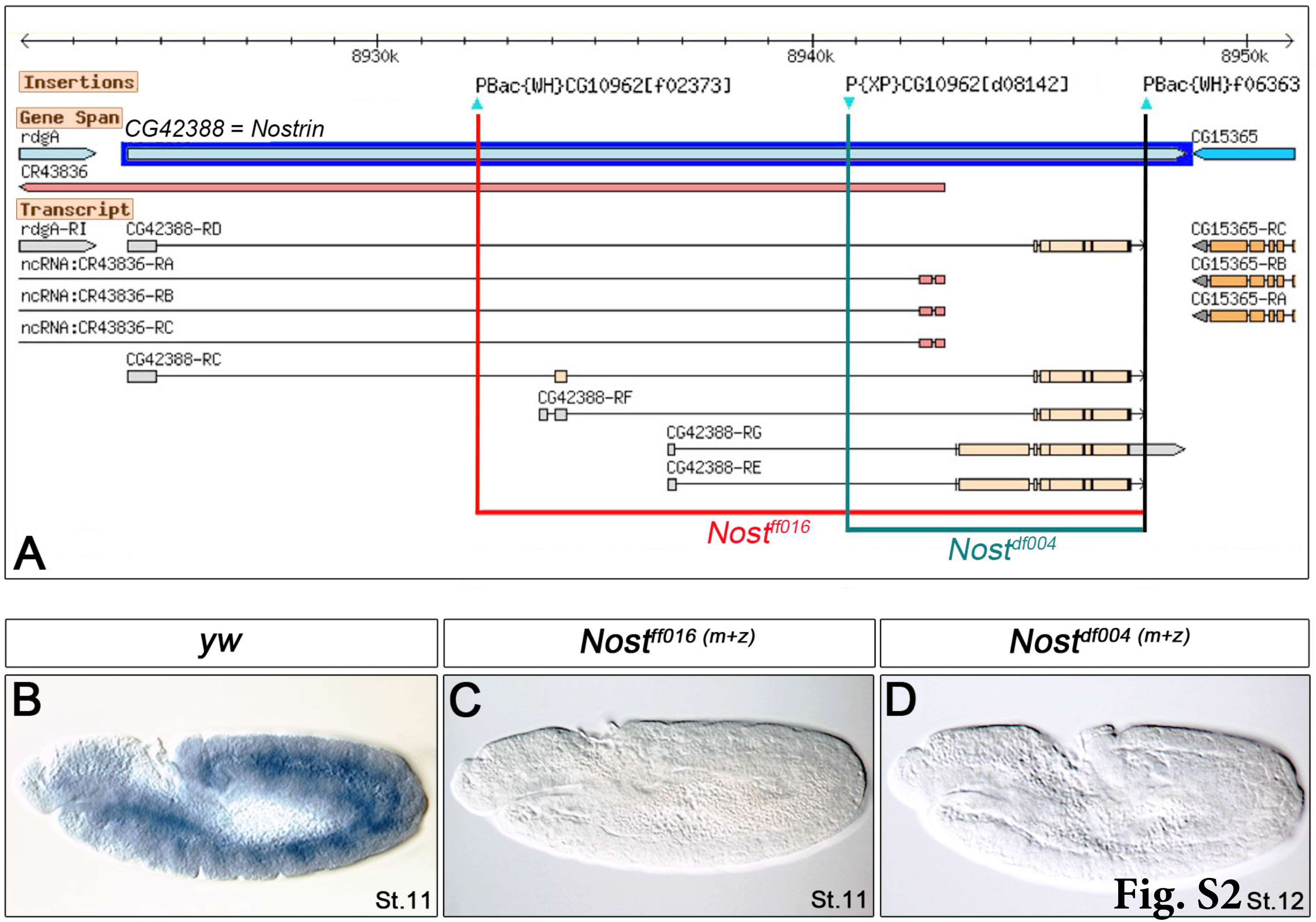
Excision mutagenesis to generate deletions at the *Drosophila Nost* locus. **(A)** Genomic map (modified from FlyBase) showing the three pBac insertions used to generate deletions in the *Nost* gene locus via recombination at their FRT sites (red bar: deletion in *Nost*^ff016^ allele; green bar: deletion in *Nost*^ddf004^ allele). **(B)** *In situ* hybridization with *Nost* antisense probe for 3’ protein coding exons in *yw* control embryo showing mesodermal expression at stage 11. **(C), (D)** *In situ* hybridizations with *Nost* antisense probe as in (B) in embryos homozygous for the two excision alleles confirms lack of *Nost* mRNA expression.

**Figure S3.**
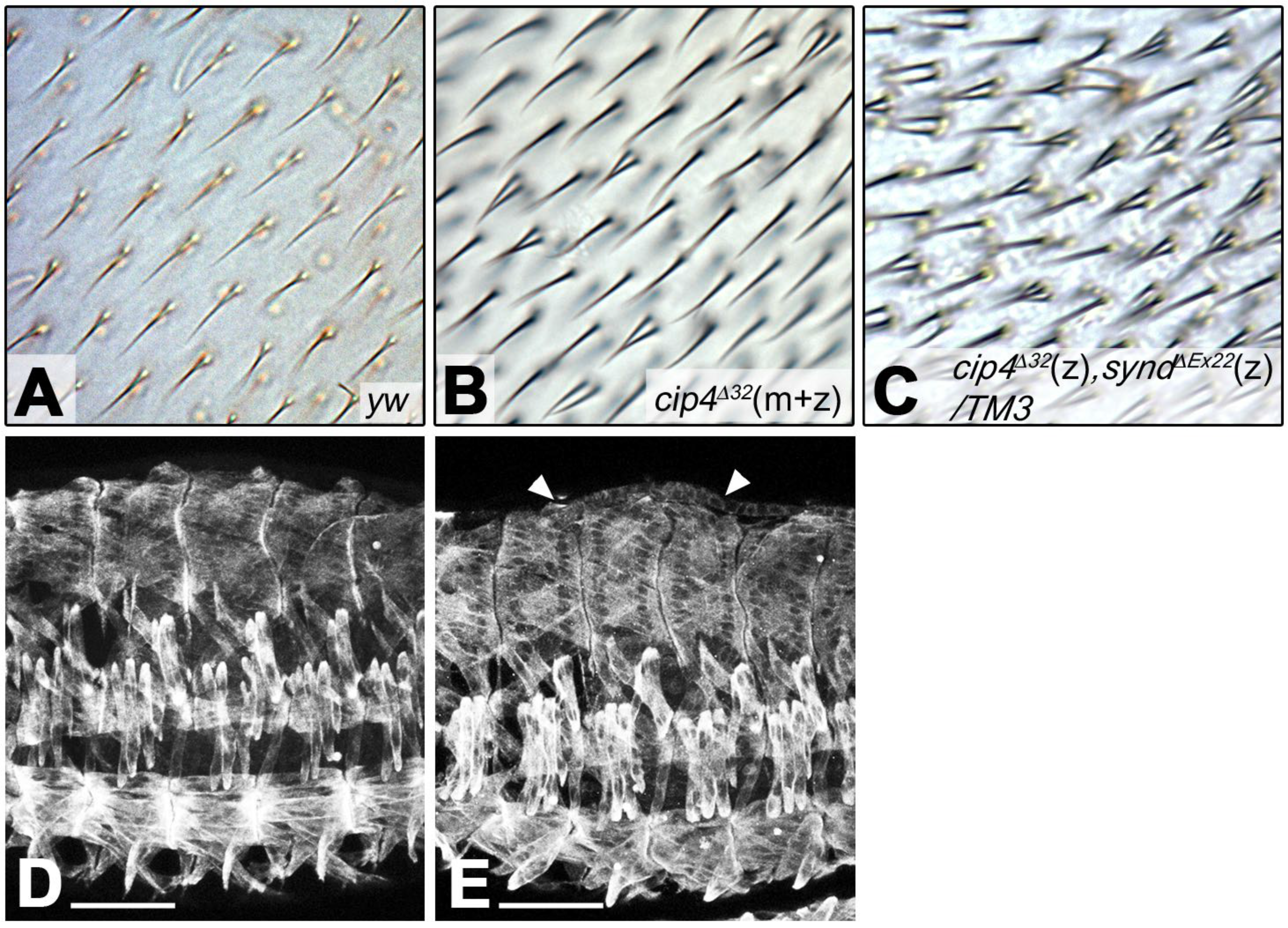
Genetic interaction of *cip4* with *synd* and phenotypes of *cip4, synd* double mutant embryos. **(A)** Single wing hairs in *yw* control wing. **(B)** Frequently duplicated wing hairs in *cip4*^Δ32^(m+z) mutant wing for comparison to (C), *cip4*^Δ32^ (z), *synd*^ΔEx22^(z)/TM3, which shows more frequently duplicated or even triplicated wing hairs in heterozygous condition of *cip synd* double mutant. **(D)** Outside view of musculature of the same stage 16 *yw* control embryo as shown in Fig. 6A (anti-Tropomyosin I). **(E)** Outside view of musculature of the same stage 16 homozygous *cip4*^Δ32^(z), *synd*^ΔEx22^(z) embryo as shown in Fig. 6B. Muscle phenotypes are not evident. A bulge at the dorsal side of the embryo (arrow heads) is caused by the expansion of the anterior midgut chamber (see Fig. 6B). Scale bars in D, E: 50μm.

**Table S1.** Somatic muscle phenotypes in *Nost*^▵f004^(m+z) *Cip4*^▵32^(m+z) double and *Nost*^▵f004^ (m+z) *Cip4*^▵32^ (m+z) *Synd*^▵Ex22^(z) triple mutant embryos.

| unfused myoblasts present | missing muscles | embryo halves, n= | frequency, % |
| --- | --- | --- | --- |
| ✓ | none | 60 | 49.6 |
| ✓ | in 1 segment | 33 | 27.3 |
| ✓ | in 2 segments | 18 | 14.9 |
| ✓ | in 3 segments | 8 | 6.6 |
| ✓ | in 4 segments | 2 | 1.6 |
| Total |  | 121 | 100 |

**B.** Myoblast fusion defects in double and triple mutants
| genotype | stage 14 | early stage 15 | late stage 15 | stage 16 |
| --- | --- | --- | --- | --- |
| **WT**<sup>1)</sup> | 4,5 ± 0,97<br>(n=30) | 7,3 ± 1,75<br>(n=30) | 10,97 ± 1,38<br>(n=30) | --- |
| **yw**<sup>2)</sup> | 4,86 ± 0,81<br>(n=35) | 7,86 ± 1,22<br>(n=125) | 10,2 ± 1,37<br>(n=125) | 11,22 ± 1,72<br>(n=95) |
| ***nost<sup>f004</sup>(m+z);;cip4<sup>32</sup>(m+z)***<sup>3)</sup> | 3,88 ± 1,11<br>(n=114) | 5,86 ± 1,98<br>(n=133) | 8,45 ± 1,97<br>(n=114) | 9,57 ± 2,85<br>(n=14) |
| ***nost<sup>f004</sup>(m+z);;cip4<sup>32</sup>(z),synd<sup>Ex22</sup>(z)***<sup>4)</sup> | 3,55 ± 1,35<br>(n=75) | 6,2 ± 1,8<br>(n=93) | 8,07 ± 1,9<br>(n=126) | 8,56 ± 2,17<br>(n=89) |

1) data from Bataillé et al., 2010; 2) own data; 3) embryos from homozygous *nost<sup>f004</sup>(m+z);;cip4<sup>32</sup>(m+z)* parents; 4) homozygous embryos from *nost<sup>f004</sup>(m+z);;cip4<sup>32</sup>(z),synd<sup>Ex22</sup>(z)/TM3, twi>>GFP* parents; ± = standard deviation; n = number of evaluated muscles, m+z, maternally plus zygotically ablated gene functions; z, only zygotically ablated gene function.

## References

Aghajanian, P., S. Takashima, M. Paul, A. Younossi-Hartenstein and V. Hartenstein, 2016 Metamorphosis of the *Drosophila* visceral musculature and its role in intestinal morphogenesis and stem cell formation. Dev Biol 420: 43–59.

Azpiazu, N., and M. Frasch, 1993 *tinman* and *bagpipe:* two homeo box genes that determine cell fates in the dorsal mesoderm of *Drosophila*. Genes Dev 7: 1325–1340.

Balagopalan, L., M. H. Chen, E. R. Geisbrecht and S. M. Abmayr, 2006 The CDM superfamily protein MBC directs myoblast fusion through a mechanism that requires phosphatidylinositol 3,4,5-triphosphate binding but is independent of direct interaction with DCrk. Mol Cell Biol 26: 9442–9455.

Bataille, L., I. Delon, J. P. Da Ponte, N. H. Brown and K. Jagla, 2010 Downstream of identity genes: muscle-type-specific regulation of the fusion process. Dev Cell 19: 317–328.

Bour, B., M. O’Brien, W. Lockwood, E. Goldstein, R. Bodmer et al., 1995 *Drosophila* MEF2, a transcription factor that is essential for myogenesis. Genes Dev. 9: 730–741.

Brinkmann, K., M. Winterhoff, S. F. Onel, J. Schultz, J. Faix et al., 2016 WHAMY is a novel actin polymerase promoting myoblast fusion, macrophage cell motility and sensory organ development in *Drosophila*. J Cell Sci 129: 604–620.

Chen, E. H., B. A. Pryce, J. A. Tzeng, G. A. Gonzalez and E. N. Olson, 2003 Control of myoblast fusion by a guanine nucleotide exchange factor, loner, and its effector ARF6. Cell 114: 751–762.

Chen, Y., J. Aardema, A. Misra and S. J. Corey, 2012 BAR proteins in cancer and blood disorders. Int J Biochem Mol Biol 3: 198–208.

Deng, S., M. Azevedo and M. Baylies, 2017 Acting on identity: Myoblast fusion and the formation of the syncytial muscle fiber. Semin Cell Dev Biol 72: 45–55.

Doherty, J. T., K. C. Lenhart, M. V. Cameron, C. P. Mack, F. L. Conlon et al., 2011 Skeletal muscle differentiation and fusion are regulated by the BAR-containing Rho-GTPase-activating protein (Rho-GAP), GRAF1. J Biol Chem 286: 25903–25921.

Dottermusch-Heidel, C., V. Groth, L. Beck and S. F. Onel, 2012 The Arf-GEF Schizo/Loner regulates N-cadherin to induce fusion competence of *Drosophila* myoblasts. Dev Biol 368: 18–27.

Duan, H., J. B. Skeath and H. T. Nguyen, 2001 *Drosophila* Lame duck, a novel member of the Gli superfamily, acts as a key regulator of myogenesis by controlling fusion-competent myoblast development. Development 128: 4489–4500.

Feng, Y., S. M. Hartig, J. E. Bechill, E. G. Blanchard, E. Caudell et al., 2010 The *Cdc42-interacting protein-4* (*CIP4*) gene knock-out mouse reveals delayed and decreased endocytosis. J Biol Chem 285: 4348–4354.

Frasch, M., M. Paddy and H. Saumweber, 1988 Developmental and mitotic behaviour of two novel groups of nuclear envelope antigens of *Drosophila melanogaster*. J. Cell Sci. 90: 247–263.

Fricke, R., C. Gohl and S. Bogdan, 2010 The F-BAR protein family Actin’ on the membrane. Commun Integr Biol 3: 89–94.

Fricke, R., C. Gohl, E. Dharmalingam, A. Grevelhorster, B. Zahedi et al., 2009 *Drosophila* Cip4/Toca-1 integrates membrane trafficking and actin dynamics through WASP and SCAR/WAVE. Curr Biol 19: 1429–1437.

Geisbrecht, E. R., S. Haralalka, S. K. Swanson, L. Florens, M. P. Washburn et al., 2008 *Drosophila* ELMO/CED-12 interacts with Myoblast city to direct myoblast fusion and ommatidial organization. Dev Biol 314: 137–149.

George, B., N. Jain, P. Fen Chong, J. Hou Tan and T. Thanabalu, 2014 Myogenesis defect due to *Toca-1* knockdown can be suppressed by expression of N-WASP. Biochim Biophys Acta 1843: 1930–1941.

Gildor, B., R. Massarwa, B. Z. Shilo and E. D. Schejter, 2009 The SCAR and WASp nucleation-promoting factors act sequentially to mediate *Drosophila* myoblast fusion. EMBO Rep 10: 1043–1050.

Hamp, J., A. Lower, C. Dottermusch-Heidel, L. Beck, B. Moussian et al., 2016 *Drosophila* Kette coordinates myoblast junction dissolution and the ratio of Scar-to-WASp during myoblast fusion. J Cell Sci 129: 3426–3436.

Haralalka, S., C. Shelton, H. N. Cartwright, F. Guo, R. Trimble et al., 2014 Live imaging provides new insights on dynamic F-actin filopodia and differential endocytosis during myoblast fusion in *Drosophila*. PLoS One 9: e114126.

Icking, A., S. Matt, N. Opitz, A. Wiesenthal, W. Muller-Esterl et al., 2005 NOSTRIN functions as a homotrimeric adaptor protein facilitating internalization of eNOS. J Cell Sci 118: 5059–5069.

Itoh, T., K. S. Erdmann, A. Roux, B. Habermann, H. Werner et al., 2005 Dynamin and the actin cytoskeleton cooperatively regulate plasma membrane invagination by BAR and F-BAR proteins. Dev Cell 9: 791–804.

Jin, P., R. Duan, F. Luo, G. Zhang, S. N. Hong et al., 2011 Competition between Blown fuse and WASP for WIP binding regulates the dynamics of WASP-dependent actin polymerization in vivo. Dev Cell 20: 623–638.

Kaipa, B. R., H. Shao, G. Schafer, T. Trinkewitz, V. Groth et al., 2013 Dock mediates Scar- and WASp-dependent actin polymerization through interaction with cell adhesion molecules in founder cells and fusion-competent myoblasts. J Cell Sci 126: 360–372.

Kessels, M. M., and B. Qualmann, 2002 Syndapins integrate N-WASP in receptor-mediated endocytosis. EMBO J 21: 6083–6094.

Kim, J. H., P. Jin, R. Duan and E. H. Chen, 2015 Mechanisms of myoblast fusion during muscle development. Curr Opin Genet Dev 32: 162–170.

Kim, S., M. Nahm, N. Kim, Y. Kwon, J. Kim et al., 2017 Graf regulates hematopoiesis through GEEC endocytosis of EGFR. Development 144: 4159–4172.

Kim, S., K. Shilagardi, S. Zhang, S. N. Hong, K. L. Sens et al., 2007 A critical function for the actin cytoskeleton in targeted exocytosis of prefusion vesicles during myoblast fusion. Dev Cell 12: 571–586.

Klapper, R., 2000 The longitudinal visceral musculature of *Drosophila melanogaster* persists through metamorphosis. Mech Dev 95: 47–54.

Knirr, S., N. Azpiazu and M. Frasch, 1999 The role of the NK-homeobox gene *slouch* (S59) in somatic muscle patterning. Development 126: 4525–4535.

Kumar, V., S. R. Alla, K. S. Krishnan and M. Ramaswami, 2009a Syndapin is dispensable for synaptic vesicle endocytosis at the *Drosophila* larval neuromuscular junction. Mol Cell Neurosci 40: 234–241.

Kumar, V., R. Fricke, D. Bhar, S. Reddy-Alla, K. S. Krishnan et al., 2009b Syndapin promotes formation of a postsynaptic membrane system in *Drosophila*. Mol Biol Cell 20: 2254–2264.

Lee, H.-H., S. Zaffran and M. Frasch, 2005 The Development of the Visceral Musculature of *Drosophila*, pp. 62–78 in Drosophila Muscle Development, edited by H. Sink. New York, Springer.

Leibfried, A., R. Fricke, M. J. Morgan, S. Bogdan and Y. Bellaiche, 2008 *Drosophila* Cip4 and WASp define a branch of the Cdc42-Par6-aPKC pathway regulating E-cadherin endocytosis. Curr Biol 18: 1639–1648.

Lenhart, K. C., A. L. Becherer, J. Li, X. Xiao, E. M. McNally et al., 2014 GRAF1 promotes ferlin-dependent myoblast fusion. Dev Biol 393: 298–311.

Lilly, B., S. Galewsky, A. Firulli, R. Schulz and E. Olson, 1994 D-MEF2: a MADS box transcription factor expressed in differentiating mesoderm and muscle cell lineages during *Drosophila* embryogenesis. Proc Natl Acad. Sci U S A 91: 5662–5666.

Liu, S., X. Xiong, X. Zhao, X. Yang and H. Wang, 2015 F-BAR family proteins, emerging regulators for cell membrane dynamic changes-from structure to human diseases. J Hematol Oncol 8: 47.

Massarwa, R., S. Carmon, B. Z. Shilo and E. D. Schejter, 2007 WIP/WASp-based actin-polymerization machinery is essential for myoblast fusion in *Drosophila*. Dev Cell 12: 557–569.

Nguyen, H., R. Bodmer, S. Abmayr, J. McDermott and N. Spoerel, 1994 *D-mef2:* a *Drosophila* mesoderm-specific MADS box-containing gene with a biphasic expression profile during embryogenesis. Proc Natl Acad Sci U S A 91: 7520–7524.

Nose, A., T. Isshiki and M. Takeichi, 1998 Regional specification of muscle progenitors in *Drosophila*: the role of the *msh* homeobox gene. Development 125: 215–223.

Önel, S. F., and R. Renkawitz-Pohl, 2009 FuRMAS: triggering myoblast fusion in *Drosophila*. Dev Dyn 238: 1513–1525.

Önel, S. F., M. B. Rust, R. Jacob and R. Renkawitz-Pohl, 2014 Tethering membrane fusion: common and different players in myoblasts and at the synapse. J Neurogenet 28: 302–315.

Parks, A. L., K. R. Cook, M. Belvin, N. A. Dompe, R. Fawcett et al., 2004 Systematic generation of high-resolution deletion coverage of the *Drosophila melanogaster* genome. Nat Genet 36: 288–292.

Patel, N. H., B. G. Condron and K. Zinn, 1994 Pair-rule expression patterns of *even-skipped* are found in both short- and long-germ beetles. Nature 367: 429–434.

Ranganayakulu, G., D. Elliott, R. Harvey and E. Olson, 1998 Divergent roles for *NK-2* class homeobox genes in cardiogenesis in flies and mice. Development 125: 3037–3048.

Richard, J. P., E. Leikina, R. Langen, W. M. Henne, M. Popova et al., 2011 Intracellular curvature-generating proteins in cell-to-cell fusion. Biochem J 440: 185–193.

Richardson, B. E., K. Beckett, S. J. Nowak and M. K. Baylies, 2007 SCAR/WAVE and Arp2/3 are crucial for cytoskeletal remodeling at the site of myoblast fusion. Development 134: 4357–4367.

Roberts-Galbraith, R. H., and K. L. Gould, 2010 Setting the F-BAR: functions and regulation of the F-BAR protein family. Cell Cycle 9: 4091–4097.

Ruiz-Gomez, M., N. Coutts, A. Price, M. Taylor and M. Bate, 2000 *Drosophila* Dumbfounded: a myoblast attractant essential for fusion. Cell 102: 189–198.

Salzer, U., J. Kostan and K. Djinovic-Carugo, 2017 Deciphering the BAR code of membrane modulators. Cell Mol Life Sci 74: 2413–2438.

Schaub, C., H. Nagaso, H. Jin and M. Frasch, 2012 *Org-1*, the *Drosophila* ortholog of Tbx1, is a direct activator of known identity genes during muscle specification. Development 139: 1001–1012.

Schmitt-Engel, C., D. Schultheis, J. Schwirz, N. Ströhlein, N. Troelenberg et al., 2015 The iBeetle large-scale RNAi screen reveals gene functions for insect development and physiology. Nat Commun 6: 7822.

Schultheis, D., M. Weißkopf, C. Schaub, S. Ansari, V. A. Dao et al., 2019 A large-scale systemic RNAi screen in the red flour beetle *Tribolium castaneum* identifies novel genes involved in arthropod muscle development. G3, submitted.

Segev, N., O. Avinoam and B. Podbilewicz, 2018 Fusogens. Curr Biol 28: R378–R380.

Sens, K. L., S. Zhang, P. Jin, R. Duan, G. Zhang et al., 2010 An invasive podosome-like structure promotes fusion pore formation during myoblast fusion. J Cell Biol 191: 1013–1027.

Sherlekar, A., and R. Rikhy, 2016 Syndapin promotes pseudocleavage furrow formation by actin organization in the syncytial *Drosophila* embryo. Mol Biol Cell 27: 20642079.

Simionescu-Bankston, A., G. Leoni, Y. Wang, P. P. Pham, A. Ramalingam et al., 2013 The N-BAR domain protein, Bin3, regulates Rac1-and Cdc42-dependent processes in myogenesis. Dev Biol 382: 160–171.

Strasburger, M., 1932 Bau, Funktion und Variabilität des Darmtractus von *Drosophila melanogaster* Meigen. Z. wiss. Zool. 140: 539–649.

Tautz, D., and C. Pfeifle, 1989 A non-radioactive *in situ* hybridization method for the localization of specific RNAs in *Drosophila* embryos reveals translational control of the segmentation gene *hunchback*. Chromosoma 98: 81–85.

Yin, Z., X.-L. Xu and M. Frasch, 1997 Regulation of the Twist target gene *tinman* by modular *cis*-regulatory elements during early mesoderm development. Development 124: 4871–4982.

Zimmermann, K., N. Opitz, J. Dedio, C. Renne, W. Muller-Esterl et al., 2002 NOSTRIN: a protein modulating nitric oxide release and subcellular distribution of endothelial nitric oxide synthase. Proc Natl Acad Sci U S A 99: 17167–17172.

Zobel, T., K. Brinkmann, N. Koch, K. Schneider, E. Seemann et al., 2015 Cooperative functions of the two F-BAR proteins Cip4 and Nostrin in the regulation of E-cadherin in epithelial morphogenesis. J Cell Sci 128: 1453.

